# Predicting the fitness costs of complex mutations

**DOI:** 10.1101/2021.03.10.434744

**Authors:** Pablo Yubero, Juan F. Poyatos

## Abstract

The fitness cost of complex pleiotropic mutations is generally difficult to assess. On the one hand, it is necessary to identify which molecular properties are directly altered by the mutation. On the other, this alteration modifies the activity of many genetic targets with uncertain consequences. Here, we examine the possibility of addressing these challenges by identifying unique predictors of these costs. To this aim, we consider mutations in the RNA polymerase (RNAP) in *Escherichia coli* as a model of complex mutations. Changes in RNAP modify the global program of transcriptional regulation, with many consequences. Among others is the difficulty to decouple the direct effect of the mutation from the response of the whole system to such mutation. A problem that we solve quantitatively with data of a set of constitutive genes, which better read the global program. We provide a statistical framework that incorporates the direct effects and other molecular variables linked to this program as predictors, which leads to the identification that some genes are more suitable predictors than others. Therefore, we not only identified which molecular properties best anticipate costs in fitness, but we also present the paradoxical result that, despite pleiotropy, specific genes serve as better predictors. These results have connotations for the understanding of the architecture of robustness in biological systems.

## INTRODUCTION

One recurrent problem in Biology is to understand the impact that mutations have on fitness (Griffiths et al. 2015). Admittedly, this topic has been the center of most recent research in Molecular Biology, with a catch. The majority of mutations, for which we have a well-defined knowledge of the underlying causes of their fitness costs, are “simple”. By simple, we refer to mutations in molecular elements with a specific function, e.g., an enzyme catalyzing a particular metabolic reaction or a transcription factor linked to the activation of a given gene.

We will not examine here fitness costs of simple mutations but alternatively of those considered “complex”. Complex mutations can be commonly established by the pleiotropic action of the molecular agents experiencing the mutation (Dudley et al. 2005). For instance, these agents could refer to a core element of the metabolic or expression cellular machinery, whose function is recognized to be highly pleiotropic. One way to further outline this definition is to add that the said molecular element is active in different contexts (He and Zhang 2005), i.e., that it presents a characteristic *environmental fitness cost map*. In this map, one represents pairs of fitness values for both the wild type (WT) and a given mutant in a set of environmental conditions (Fig. 1A). Impairment of a pleiotropic agent should lead to a proportional decrease in fitness characterized by a global scale factor compared to simple mutations that uniquely display fitness costs in specific situations (Fig. 1B).

**Figure 1.**
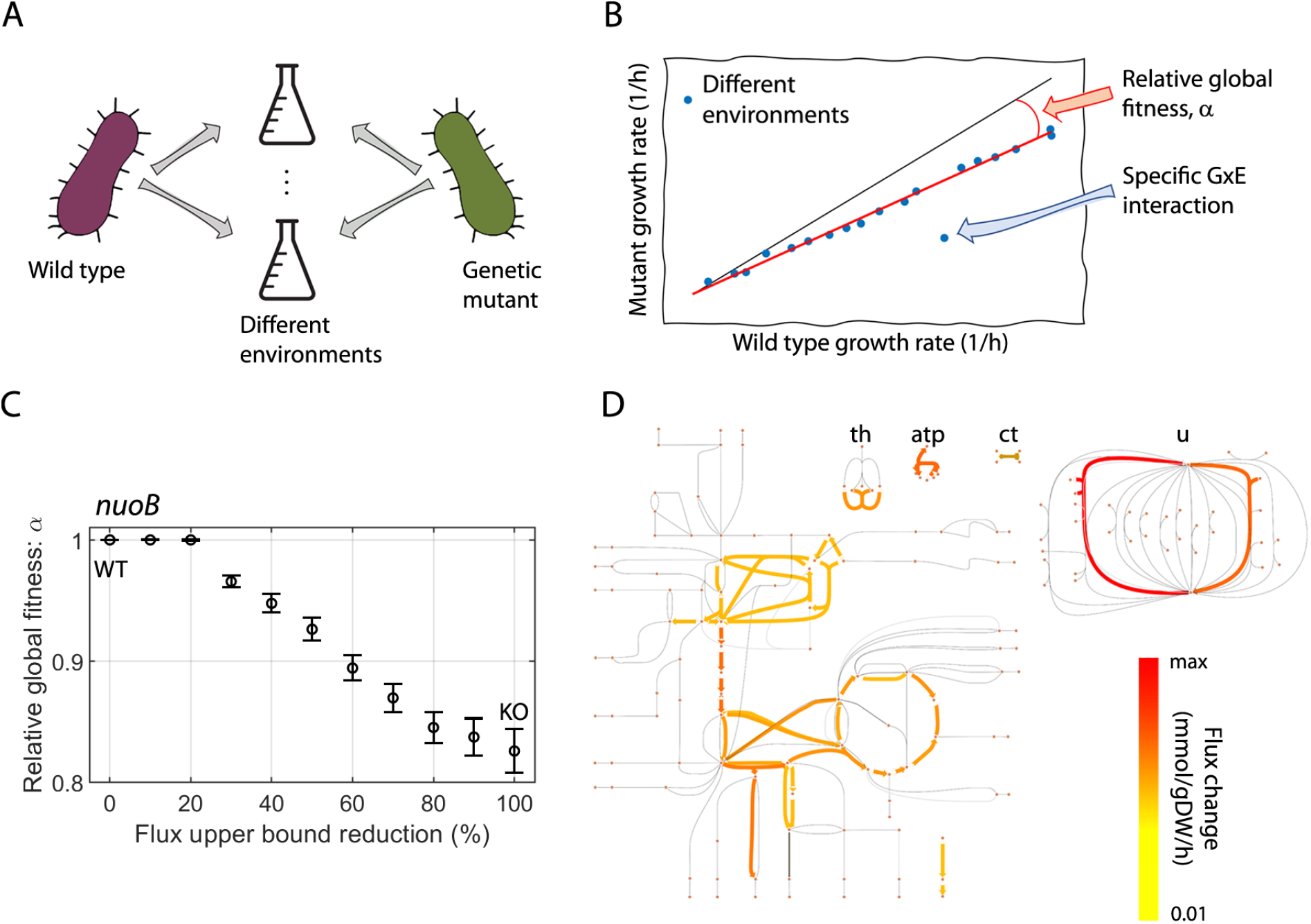
Complex mutations have a characteristic *environmental fitness cost map* because they affect globally. (**A**) Environmental fitness cost maps are obtained by measuring, and comparing, the phenotype of a genetic mutant and its WT relative in different environments. In our case, we focus on growth rate. (**B**) Sketch of an environmental fitness cost map. It facilitates the identification of complex mutations and specific gene-environment interactions (GxE). While the former is a rescaling of the fitness in most environments (red line, relative global fitness *α*), the latter are shown as outliers from this trend. (**C**) We computed the value of *α* for multiple mutants of *nuoB* using a computational metabolic model of *E. coli* (Methods). Error bars denote the 95% CI of the slope after robustly fitting data to a linear trend (as in panel B; Methods; 100% flux reduction denotes a knockout, KO). (**D**) Sketch of *E. coli*’s *nuoB* KO metabolism with the median flux change across all environments. We show only the 10% of reactions that are most affected by the mutation. Transhydrogenase (th), ATP synthase (atp), carbonate (ct), and ubiquinone reduction/oxidation (u) pathways are also shown.

In this work, we initially exemplify these concepts using a genome-wide computational model of *Escherichia coli*’s metabolism (Feist et al. 2007). We then consider the RNA polymerase (RNAP) as experimental model. Three different mutations of the gene *rpoB*, which encodes the *f3* subunit of the RNAP, follow the characteristic environmental fitness cost map of a complex mutation. Indeed, mutations in *rpoB*, usually obtained in response to rifamycins (Rif) (Goldstein 2014) –a class of antibiotics–, have been studied in many species and they entail a long list of pleiotropic effects (Jin and Gross 1989; Tóth et al. 2003; Cai et al. 2017; Karthik et al. 2019).

Once we define these mutations as complex, we then ask what set of molecular properties could be *a priori* relevant to understand their cost in fitness. We thus hypothesize several features, which organize in two broad categories, linked to the global program of transcription and the alarmone (p)ppGpp, or ppGpp onwards.

The former is motivated by the ubiquitous role of the RNAP in gene expression and its coupling to the growth rate. In fact, early works attributed fitness costs to a decreased transcriptional efficiency of the RNAP in *E. coli* (Reynolds 2000), while subsequent studies found larger, genome-wide, transcriptional reprogramming in *Pseudomonas Aeruginosa* (Qi et al. 2014), *Mycobacterium Tuberculosis* (Trauner et al. 2018) and *E. coli* (Wytock et al. 2020) that was not clearly connected to these costs. Our work will enable us to reexamine these issues.

The second broad category includes different features of the interaction between the RNAP and ppGpp mediated by the gene *dksA* (Paul et al. 2004; Irving and Corrigan 2018; Sanchez-Vazquez et al. 2019). Notably, the RNAP associated with *rpoB* mutants was found to work like a stringent RNAP (Zhou and Jin 1998), and an altered stringent response was held responsible for fitness costs in *E. coli* (Wytock et al. 2020). On top of all, the concentration of ppGpp tightly controls optimal resource allocation and hence, growth rate (Zhu and Dai 2019).

Finally, we quantify all these properties in a collection of constitutive genes as “reporters”. These genes are useful for reading the RNAP regulatory signal since they do not present any class of specific regulation (Schaechter et al. 1958; Maaløe 1979).

Armed with this data collection, we develop a quantitative framework to predict fitness costs. This leads us to reconsider earlier results. Transcriptional efficiency, i.e., the rate of mRNA production does emerge as a relevant determinant. However, comparing transcription levels between a WT and a mutant that grows at a slower rate calls for special care. Indeed, empirical laws of resource allocation show that gene expression in general, and transcriptional promoter activity in particular, are structurally dependent on the availability of global resources, which in turn, impact growth rate (Liang et al. 1999; Klumpp and Hwa 2008; Klumpp et al. 2009). This is all captured in our results.

Note that while in this example we had some knowledge of the biology involved, in general, our approach does not necessarily need a mechanistic rationale to select a particular predictor. And, although this could seem a significant drawback, it can, in turn, serve to guide research in situations where the origin of fitness costs is unknown. The statistical model can potentially integrate any number of predictors without prior knowledge about their relevance. In such a case, however, the number of experimental points needed to distinguish spurious correlations from significant ones would quickly increase. These are common problems, of course, in other theoretical and applied areas where multiple regression analysis is applied, e.g., Quantitative Genetics (Falconer and Mackay 1996) or Ecology (Johnson and Omland 2004).

More broadly, our work contributes to the general program of predicting cellular phenotypes from a molecular basis by effectively decreasing the dimensionality assumed to determine such phenotypes and has implications for our comprehension of the architecture of robustness in biological systems.

## RESULTS

### Complex mutations display global fitness costs

We first explore complex mutations *in silico*, using a genome-scale metabolic model. Specifically, we employed one convenient model of *Escherichia coli* that incorporates 1260 open reading frames (ORFs) and 2077 reactions (Feist et al. 2007). We simulate the effect of a mutation on a given enzyme by constraining the fluxes of the reactions in which it participates. Then, we compute the fitness of the WT and mutant strains in a minimal medium supplemented with one of 174 carbon sources (Fig. 1A, Methods). This enables us to distinguish between a global effect of the mutation, and specific gene-environment interactions through the environmental fitness cost map (Fig. 1B).

Enzymes involved in the energetic regulation of the metabolism are potential candidates for complex mutations. As a case study, we examined a series of *nuoB* mutants, an oxidoreductase which is part of the respiratory chain, that spanned the entire range of the effect of a mutation: from unconstrained (WT) to null (knockout) flux. Figure 1C indicates that mutants manifest a stronger decrease in *relative global fitness* (*α*; *α*<1 indicating fitness costs) for larger effects of the mutation. In the limiting case, when the reactions are turned off, we obtain the relative global fitness of the *nuoB* knockout (83%). Note that the complex character of these mutations is linked to a considerable reorganization of metabolic fluxes (Fig. 1D; see Supplementary Text and Fig. S1 for further examples and a comprehensive discussion of these mutations).

Overall, complex mutations manifest themselves in a multitude of different environments and are not specific to a particular external cue. This highlights the broader reach of these mutations and their coupling to core enzymes involved in cell growth.

### Mutations in *rpoB* are complex

We next establish how RNAP mutants represent a well-grounded experimental model system for complex mutations given RNAP’s essential role during gene expression and cellular growth. Specifically, we consider the WT strain REL606 of the bacterium *E. coli* (Barrick et al. 2009) and three mutant derivatives in the *rpoB* gene (with the following amino acid substitutions: H526L, S512Y, and Q513P) that have been selected experimentally through Rif resistance (Garibyan et al. 2003; Jin and Gross 1988).

To obtain an experimental environmental fitness cost map, we measured the growth rate of the four strains in M9 minimal media with different carbon sources (Methods). Figure 2 shows this map for the three mutants. We observe that while the derivative H526L (Fig. 2A) exhibits no fitness cost, S512Y (Fig. 2B) and Q513P (Fig. 2C) exhibit mild 4% and large 24% costs, respectively. This global response is similar to the one produced by complex mutations in the genome-scale metabolic model in the previous section. In this case, since these mutations correspond to RNAP (localized in the *rpoB* gene), we can characterize a set of molecular features directly related to the change in transcriptional performance. Ultimately, we will assess these features as potential candidates for anticipating the fitness cost of complex mutations in a statistical model.

**Figure 2.**
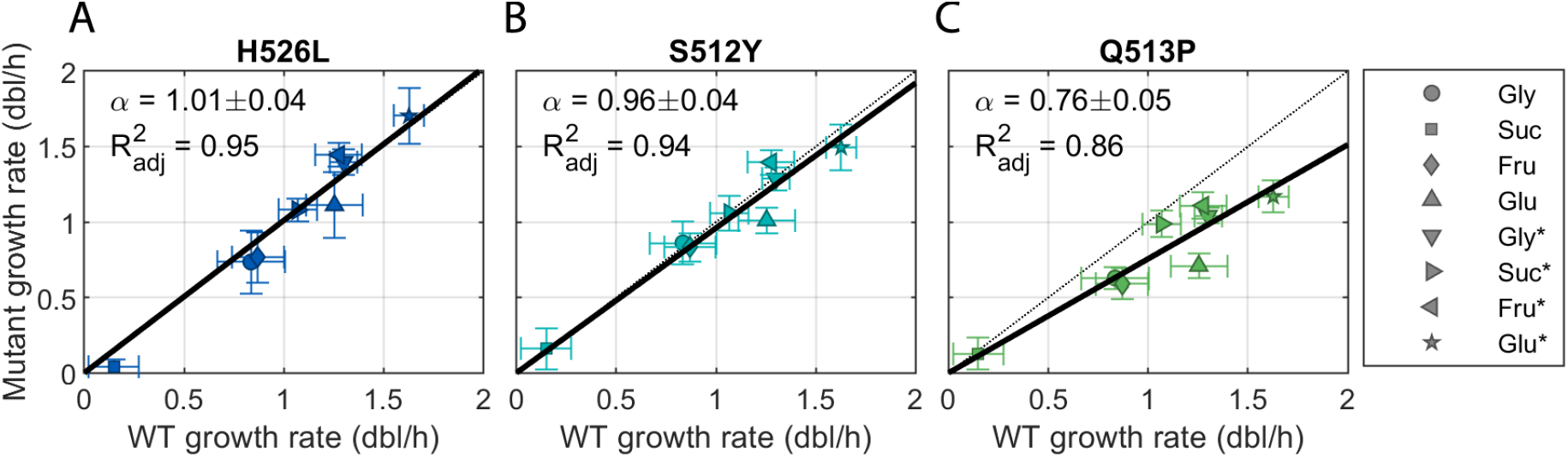
Experimental *rpoB* mutants display global fitness costs. (**A-C**) Growth rate of the three *rpoB* mutant strains (H526L, S512Y, and Q513P) and their WT relative in eight different growth media (markers, asterisks denote the addition of casamino acids; Methods). Their fitness is proportional to that of the WT, and hence can be described by their relative global fitness *α* (with its 95% CI interval). Error bars denote one standard deviation among 12 replicates.

### Mutations in *rpoB* alter the global transcriptional program

We quantified changes in the transcriptional activity of the RNAP by measuring the promoter activity (PA), i.e., the rate of mRNA production. As gene expression is strongly dependent on the growth rate, and consequently on the availability of global resources (Liang et al. 1999; Klumpp et al. 2009), changes in PA observed in the mutants present two possible causes. One is associated to a decrease in growth rate, while a second is directly linked to changes in the functional activity of the mutant RNAP (Utrilla et al. 2016). To uncouple these effects, we measured PA as a function of growth rate during balanced growth in multiple carbon sources (Fig. 3A). We introduced the notion of the *total* and *direct* promoter activity changes PA_*T*_ and PA_*D*_, respectively (Fig. 3B). While PA_*T*_ measures the difference in PA between the WT and a mutant in a given condition (and different growth rates), PA_*D*_ is the expected change in PA between the WT and a mutant when growing at the same rate (and different environmental conditions). This second measure quantifies in this way the potential change in the activity of the mutated RNAP controlling for changes in the availability of global resources due to fitness costs.

**Figure 3.**
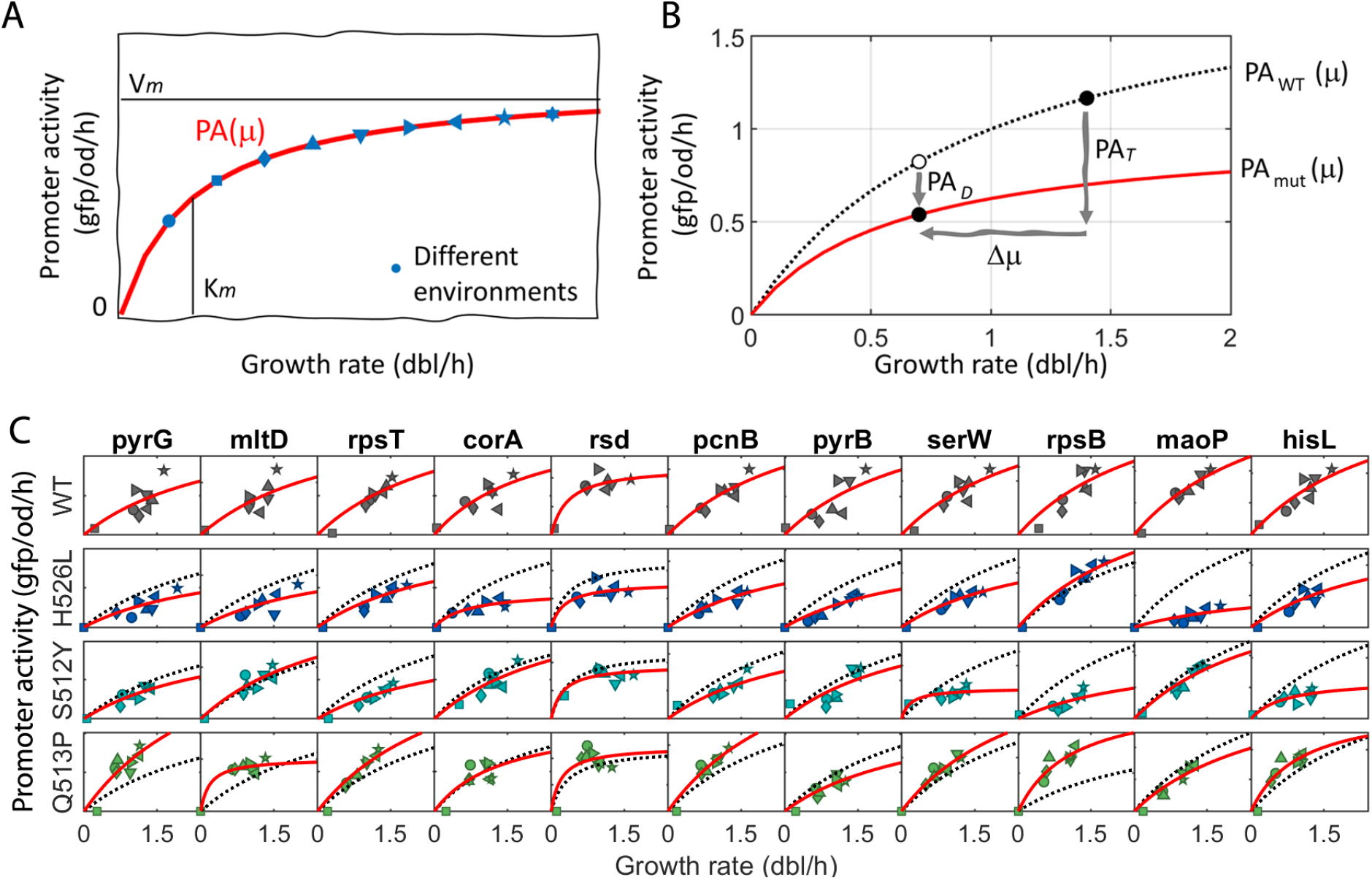
Uncoupling the total and direct effects of mutations on promoter activity. (**A**) Sketch illustrating the typical growth-rate dependency of the promoter activity of constitutive genes (red line). These are obtained from PA and growth-rate values during balanced growth in media with different carbon sources (blue symbols) and they are characterized by V_*m*_ (the maximal PA), and K_*m*_ (growth rate at which PA is half-maximal). (**B**) Sketch depicting the difference between the total and direct effects of a mutation, PA_*T*_, and PA_*D*_ respectively. PA_*T*_ measures the change in PA between the WT and the mutant in the same environment (black solid circles) but at different growth rates due to fitness costs (Δ*μ*). Quantifying the PA(*μ*) profiles in the WT and mutant (black dotted, and red solid lines respectively) enables us to capture PA_*D*_, which measures the expected change in PA when WT and mutant grow at the same rate. (**C**) PA(*μ*) profiles (red lines) of eleven constitutive genes in an array of growth media in all four strains (markers and colors, respectively, as in Fig. 2). The corresponding profile of the WT is also shown for comparability (black dotted line).

We experimentally measure PA in all strains as the accumulation rate of a reporter green fluorescent protein (GFP) of a selected set of promoters (Methods). We selected eleven *constitutive* promoters available in a reporter plasmid library (Zaslaver et al. 2006). Constitutive genes are particularly suitable because their expression does not rely on the concentration of any specific transcription factor, and thus they read the availability of global resources and the performance of the pool of RNAPs (Schaechter et al. 1958; Maaløe 1979; Klumpp and Hwa 2008). We then model the growth-rate dependencies of promoter activities, PA(*μ*), from PA measurements during exponential growth in eight different media (Methods).

Figure 3C shows the growth-rate dependencies of the promoter activities of the selected genes, in all strains, together with the best fit to a Michaelis-Menten equation PA(*μ*) = *V*_*m*_*μ* /(*K*_*m*_ + *μ*) with parameters V_*m*_, maximum expression, and K_*m*_, growth rate at which PA is half-maximal (Fig. 3A, Methods). We recovered not only that, in general, each promoter follows a specific profile with different parameters V_*m*_ and K_*m*_, but also that some of them reside in the linear regime with large K_*m*_ (Liang et al. 1999; Gerosa et al. 2013; Yubero and Poyatos 2020).

Most importantly, the activity of promoters in the RNAP mutant strains still follow hyperbolic patterns although different across strains. We found a significant tendency of H526L and S512Y towards smaller values of V_*m*_ whereas Q513P displayed a general decrease in K_*m*_ (Fig. S2A-B). However, the quantitative change in these parameters are mutation- and promoter-specific. Therefore, changes in these profiles, i.e., in the global transcriptional program, are candidates for predictors of fitness costs.

The availability of a predictive model of PA(*μ*) for all promoters in all strains enables us to distinguish between the direct effect of a mutation, PA_*D*_ to the total change in promoter activity PA_*T*_. Interestingly, in most promoters, we observe significant direct effects. Even if RNAP mutations do not produce fitness costs, as in strain H526L, most promoter activities are significantly altered in a consistent manner across environments (*>* 80%, Fig. S2C-H). Hence, apart from the total effects on PA, we also consider separately the direct effects as potential fitness predictors.

### Mutations in *rpoB* alter the action of ppGpp-RNAP

The performance of the RNAP is strongly dependent on its interaction with the alarmone ppGpp playing a pivotal role in controlling growth rate in both minimal and rich media (Irving and Corrigan 2018; Potrykus et al. 2011; Zhu and Dai 2019) and during the stationary phase (Hirsch and Elliott 2002). Besides, changes in the concentration of ppGpp, together with the presence of *dksA*, alters the genome-wide transcriptional pattern with profound consequences in resource allocation (Paul et al. 2004; Zhu and Dai 2019; Sanchez-Vazquez et al. 2019). Since some *rpoB* mutants also display defective RNAP-ppGpp action (Zhou and Jin 1998), we posit that mutations should also impact both growth and transcription during the stringent response at the exit of the exponential phase, and during the stationary phase.

Thus, we considered the following three proxies to quantitatively assess alterations in RNAP-ppGpp interactions: the promoter activity and protein level during stationary phase and the deceleration in growth rate during the stringent response. The first assesses the transcriptional reprogramming in stationary phase. The second is a measure of the aggregate effect of PA deregulation during both balanced growth and stationary phase. Finally, the deceleration rate measures the efficiency of RNAP-ppGpp in arresting growth.

Firstly, we measured the promoter activities in stationary phase during the last two hours of the experiment (PA_*f*_, Fig. 4A; note that other time windows produce qualitatively similar results). This parameter describes the appropriate ability of the pair RNAP-ppGpp to reprogram transcription when nutrients are depleted. We observe that only a subset of 4, 1, and 3 promoters in strains H526L, S512Y and Q513P, respectively, have a significant under/over-activity in stationary phase across different growth media.

**Figure 4.**
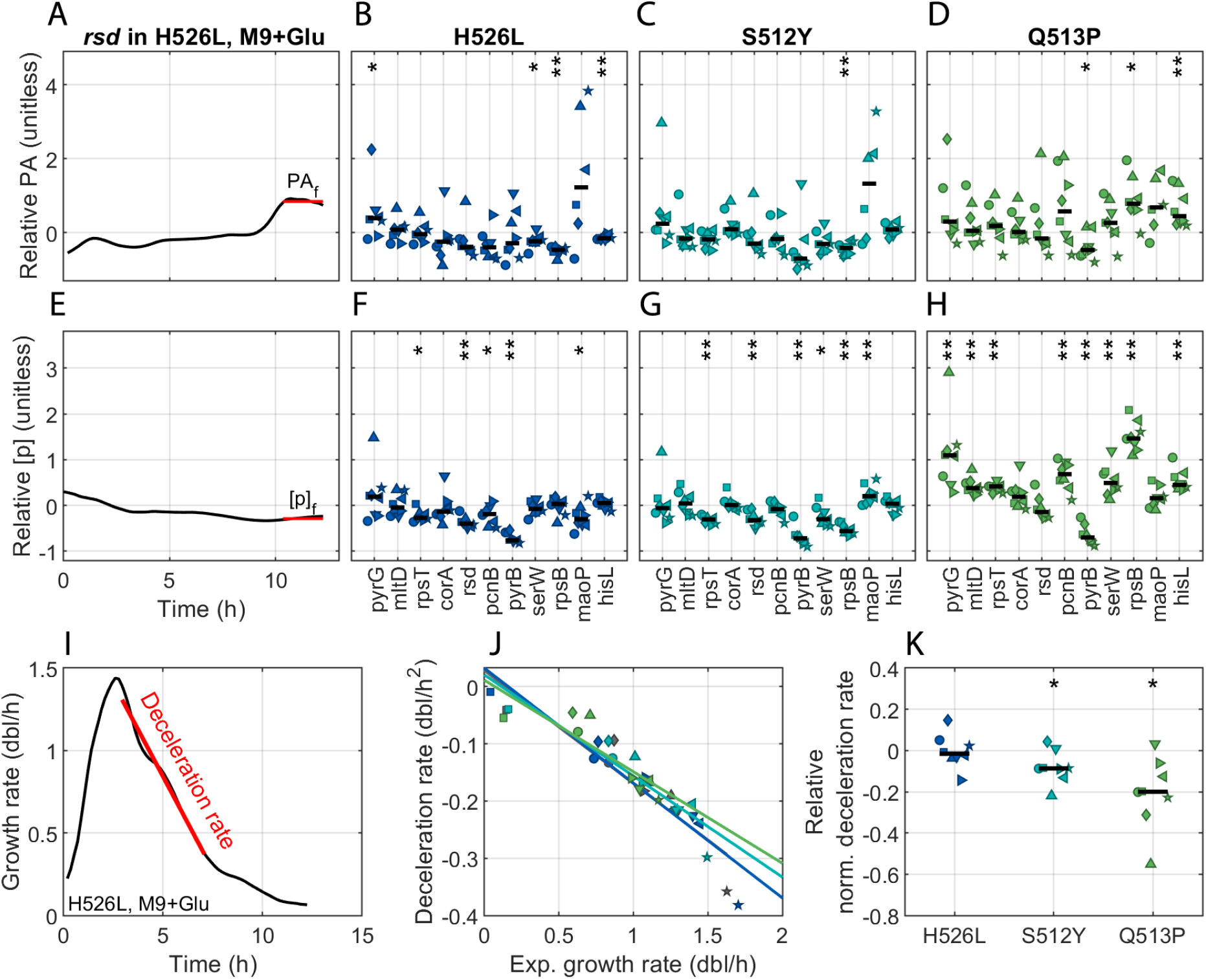
Promoter activity and protein concentration during stationary phase, and the deceleration rate constitute additional potential predictors of fitness costs. (**A**) Relative final promoter activity (ratio mutant PA_*f*_ over WT) of the *rsd* gene. (**B-D**) Relative PA_*f*_ of all promoters and mutant backgrounds (x-axis). (**E**) Relative final protein level (ratio mutant [p]_*f*_ over WT) of *rsd*. (**F-H**) Relative [p]_*f*_ of all promoters and mutant backgrounds (x-axis). **I**) deceleration rate during growth in M9 and glucose of H526L. (**J**) Deceleration rates correlate strongly with the exponential growth rates reached in that particular media (markers) in all strains (colors; Pearson’s *ρ <* − 0.86 and p<0.01 in all strains). (**K**) Even when controlling for this correlation, the relative deceleration rates of different mutants differ significantly. Note that while the first two scores are measured during the last two hours of the experiment when cultures are in stationary phase (red horizontal lines), the deceleration rate is computed from the change in growth rate right after the exponential phase (slope of the red line). In all panels, we tested a homogeneous response, either positive or negative, across all environments using a two-sided Wilcoxon sign rank test for medians (* p<0.05 and ** p<0.01). Colors and markers denote strain and media composition as in Fig.2.

Secondly, in analogy to PA_*f*_, we measure the protein level also in stationary phase (p_*f*_, Fig. 4E) to assess the combined effect of reduced promoter activity and growth rate. We find that these are more often altered than PA_*f*_, although the responses are still mutation- and promoter-specific (Fig. 4F-H). Note that p_*f*_ relative to the WT, tend to be negative in strains H526L and S512Y as opposed to Q513P.

Finally, we used the deceleration rate as a proxy of the interaction RNAP-ppGpp at the onset of the stringent response, given its fundamental role in arresting growth at the exit of the exponential phase. We measure the deceleration rate as the slope of the linear fit to the instantaneous growth rate during 4h after the exponential phase (Fig. 4I, again, other time windows produce similar results). Unsurprisingly, across all strains we observed a strong negative linear correlation between the deceleration rate and the growth rate during balanced growth (Fig. 4J). Thus, reaching a larger growth rate during exponential phase leads to a faster deceleration rate during growth arrest. Then, we searched for changes in the *normalized* deceleration rates across mutants, which controls for the respective exponential phase growth rates. Figure 4K shows that both strains with fitness costs display a significantly reduced normalized deceleration rate with respect to the WT across environments.

### A statistical model for complex fitness predictions

The characterization of all previous features equipped us with the necessary data to introduce a statistical model capable of explaining the fitness costs of three *rpoB* mutants in eight different growth media from specific molecular determinants. Given the uncoordinated changes in expression observed in the previous sections, not only do we seek which determinants are best suited for fitness costs prediction but also of which reporter genes.

Specifically, we considered the following predictors related to gene expression: the total and direct promoter activity changes PA_*T*_ and PA_*D*_, respectively; the global transcriptional program parameters V_*m*_ and K_*m*_; the promoter activity during stationary phase PA_*f*_; the protein level during stationary phase *p*_*f*_, and the normalized deceleration rate during growth arrest *∂*_*tμ*_. The model describes the relative growth rate of mutants as a function of the relative change of predictors. The expression for each gene, in Wilkinson notation, is:

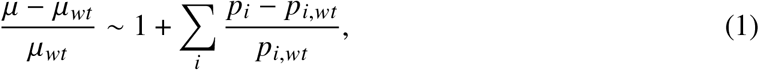

where *μ* is the growth rate; *p*_*i*_ is the i-eth predictor; the subscript *wt* denotes the WT strain and 1 refers to a constant intercept. Therefore, a positive parameter estimate implies that the relative change of the predictor correlates positively with the relative change in the growth of the mutant (Fig. S3 shows all cross-correlations between variables). Each model integrates data of the three mutants during growth in the eight media, fitting a total of 24 points.

With the statistical model, we seek which genes best describe the fitness changes and with which combinations of predictors. To do so, we used an algorithm with a step-wise addition and subtraction of predictors to an initially constant model following Bayes’ information criterion to prevent overfitting. Figure 5 and Table 1 show the results, where we observe that all promoters reach a convenient root mean squared error (RMSE) and 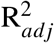 (Fig. 5A), with the exception of *corA*, an ion transporter; *pyrB*, part of the pyrimidine biosynthesis pathway; and *pcnB* involved in RNA polyadenylation.

**Table 1.** Linear models for the anticipation of fitness costs. We show the coefficients of the predictors (columns) obtained for the data set of each promoter (rows). The number in parentheses is the standard error of the coefficient in the last decimal digit shown. The last column contains the root mean squared errors as a measure of goodness of fit. Models were selected in a step-wise manner following the Bayesian information criterion (Methods). † genes with largest rmse that fit poorly the fitness costs.

| | (Intercept) | PA<sub>T</sub> | PA<sub>D</sub> | V<sub>m</sub> | K<sub>m</sub> | PA<sub>f</sub> | p<sub>f</sub> | $\partial_t \mu$ | RMSE |
| --- | --- | --- | --- | --- | --- | --- | --- | --- | --- |
| *hisL* | 0.00(6) | 0.32(8) | -0.54(8) | 0.3(1) | - | - | -0.11(5) | - | 0.096 |
| *rsd* | -1.6(4) | 0.85(4) | -0.60(4) | -2.3(5) | -3.91(4) | - | - | - | 0.114 |
| *serW* | -0.13(4) | 0.32(8) | -0.3(1) | - | - | - | -0.26(5) | - | 0.146 |
| *rpsT* | -0.09(3) | - | - | - | - | - | -0.31(6) | - | 0.149 |
| *maoP* | -0.18(7) | - | -0.07(3) | -0.3(2) | - | - | - | -0.19(4) | 0.188 |
| *rpsB* | -0.11(5) | - | - | 0.4(1) | - | - | -0.19(3) | - | 0.189 |
| *mltD* | -0.10(5) | 0.5(1) | -0.6(1) | - | - | - | - | - | 0.225 |
| *pyrG* | -0.11(6) | 0.4(1) | -0.5(1) | - | - | - | -0.10(6) | - | 0.230 |
| *corA*<sup>†</sup> | -0.5(3) | - | -3.0(5) | - | 1.7(4) | -1.8(4) | 3.7(6) | -2.1(7) | 0.933 |
| *pyrB*<sup>†</sup> | 1.7(6) | - | - | - | - | - | 2.1(9) | - | 1.75 |
| *pcnB*<sup>†</sup> | 0.5(4) | - | - | - | - | - | - | - | 1.89 |
| Median | -0.10 | 0.40 | -0.54 | 0 | -1.1 | -1.8 | -0.11 | -1.17 | 0.189 |

**Figure 5.**
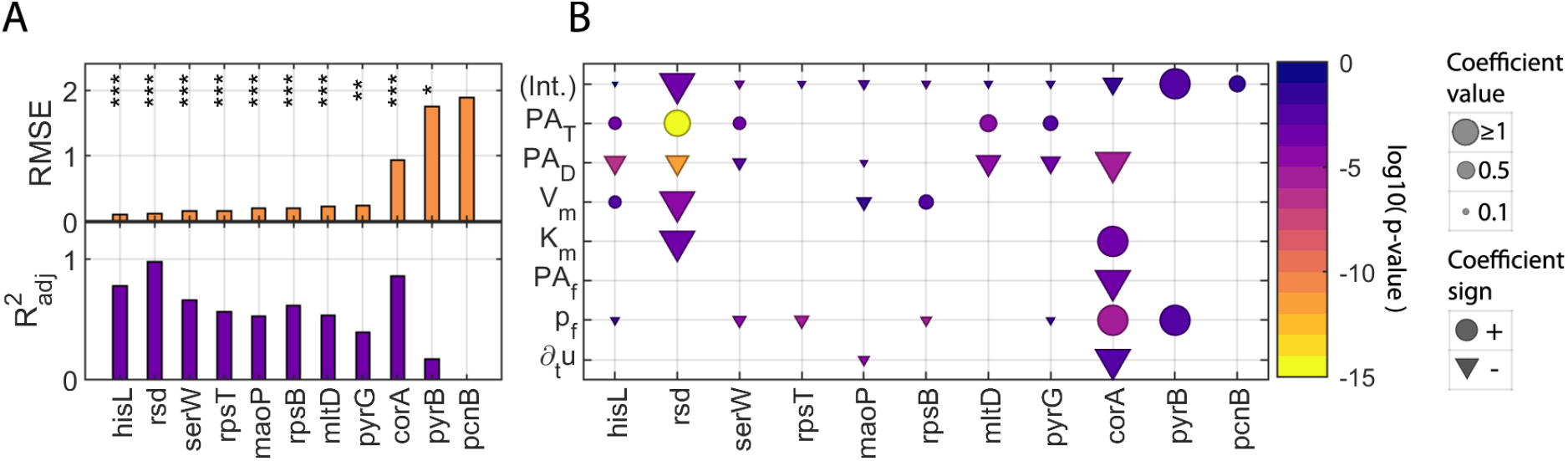
Anticipating fitness costs from molecular predictors of a variety of promoters. To predict the fitness costs of three *rpoB* mutants growing in eight different media conditions, we used a linear model with step-wise addition and subtraction of predictors following the Bayesian information criterion to avoid overfitting. (**A**) Goodness of fit as described by the root mean squared error (RMSE, top) and the adjusted *R*^2^ (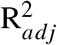 bottom) of the final linear models (* p<0.05, ** p<0.01 and *** p<0.001). (**B**) Model coefficient values (size, clipped to 1 for comparability, see Table 1), sign (marker), and significance (t-test p-value; colors are proportional to its log_10_) of each predictor and each promoter in the final linear models.

Moreover, the structure of the best models for each gene is represented in Fig. 5B. We observe a clear pattern of PA_*T*_ and PA_*D*_ as the principal predictors of fitness costs. Interestingly, the coefficients for PA_*D*_ and PA_*T*_ have opposite signs across all promoters studied, likely highlighting a general mechanism. On the one hand, a positive coefficient of PA_*T*_ implies that mutations in *rpoB* preserve the general shape of PA(*μ*) profiles as a monotonically increasing function (Fig. S4A). On the other hand, a negative coefficient of PA_*D*_ highlights that for a fixed growth rate, larger fitness costs are associated with the overexpression of constitutive promoters (Fig. S4B). This effect is clear when observing the PA(*μ*) profiles of the strain with the largest fitness cost (Q513P in Fig. 3C).

Overall, we find that a multivariate regression with as little as four (median) predictors anticipates the fitness costs of different *rpoB* mutants growing in a variety of carbon sources.

## DISCUSSION

One encounters three potential problems when characterizing the fitness costs of complex mutations: 1/to define which molecular elements are likely subjects of complex mutations, 2/to recognize which of the molecular features altered by these mutations are driving the costs, and 3/to identify whether some specific target elements (of the molecular agent) can act as a distinctive reporter of such modified features and, in this way, of the costs. We find an answer to the first problem with the use of the environmental fitness cost map and to the second by dissecting a set of potential predictors, quantified in reporter genes, that are ultimately integrated into a statistical model. By identifying patterns in the models of a variety of genes, this approach also helps us to resolve the third problem: which targets could be most relevant to predict the fitness costs of mutations.

That we observe complex mutations in a metabolic model supports the idea that they are likely prevalent in regulatory networks and hence, in biological systems. Moreover, we verify that such perturbations are associated with fundamental organismal functions and a larger system-level reprogramming as they are apparent in all environments. The larger reach of these mutations could be connected to pleiotropic effects. Here we find not only that mutations in *E. coli*’s RNAP are complex, but also that their phenotype changes are highly specific to the mutation.

The use of RNAP as an experimental (model) system presents some advantages. First, we can select predictors with clear biological significance. These predictors are mainly related to either the performance of RNAP or its interaction with the alarmone ppGpp. Second, we can test the validity of our approach to earlier discussions on the fitness costs of RNAP mutations. Last, we can consider constitutive genes as an appropriate set of reporters. These genes are valid reporters of both direct effects on transcriptional efficiency and indirect ones on cell physiology (see below). Given that the sensitivity to the latter (the global program of transcription) is gene-dependent (Liang et al. 1999; Gerosa et al. 2013; Yubero and Poyatos 2020), we identify some genes within this class that are eventually better predictors than others through the same subset of variables to acceptable levels (but three genes fail terribly in the task; see Fig. S5 for analysis of specific molecular attributes).

We propose that genes that perform better are somehow sensitive to growth rate. This sensitivity could be read through the changes in a set of features, e.g., their expression, as in the following scenarios. First, a gene whose expression is highly robust to a mutation producing fitness costs will likely fail at predicting these costs, as even in the presence of such mutation there will be no observable change in the features. Second, a gene that is disrupted by the presence of the mutation will again be a bad predictor as its expression becomes irrelevant or unreliable. We hypothesize that in between these scenarios, there are a few genes whose predictability is maximal as they are only partially affected by the mutation. We verify this by quantifying the overall effect of a mutation on a gene as the sum of the squared relative change of the predictors included in the statistical framework (Fig.S6).

Moreover, a comparison of the expression response of a mutant to the WT for a *fixed* growth rate could further confirm constitutive genes as the best reporters of fitness costs. To this aim, we used RNA-seq data of the *rpoB* mutant E546V and its WT ancestor (Utrilla et al. 2016). The transcriptional changes produced by E546V at two different (fixed) growth rates correlate only slightly (Spearman’s *ρ* = 0.12; Fig. S7A). But most importantly, we found that this correlation greatly originates from the response of constitutive rather than regulated genes (Fig. S7B). Should this be a general case, it highlights not only that the transcriptional changes produced by a mutation in *rpoB* are dependent on the growth rate, but also that constitutive genes display a more coordinated response. Consequently, these genes are probable better fitness costs predictors than genes subjected to more specific regulation. In other words, the regulatory network can partially buffer the transcriptional changes produced by the mutant RNAP.

### Specific implications to the interplay between transcriptional efficiency and fitness cost in Rifresistant *rpoB* mutants

Mutations in *rpoB* are most commonly found in antibiotic resistance and adaptive evolution experiments and have been studied extensively due to their implications in tuning fitness. More specifically, mutants producing fitness costs have been traditionally correlated to changes in the transcriptional efficiency of the mutant RNAPs. However, there are several issues with the previous studies.

First, changes in transcriptional efficiency are promoter, environment, and (mutant) strain-dependent. A restricted number of any of these variables limits, therefore, the generality of these results. However, alleviating this largely increases the cost and difficulties of such studies. Our data set is a compromise that allows having a broader view of the impact of mutations in *rpoB* on the transcription of different promoters across multiple growth media.

Second, there exists a core dependency between growth rate and gene expression unaccounted for in previous studies. This relationship is most evident in the PA(*μ*) profiles of constitutive genes as PA increases together with growth rate (Fig. 3) what anticipates a decrease in transcription when cells grow at a reduced rate even in the absence of mutations. Moreover, mutations in *rpoB* directly affect the transcriptional activity of the RNAP producing fitness costs, which in turn, further constrain the efficiency of the RNAP.

For this reason, total changes in PA have a direct contribution to the mutation, and what we called an *indirect* contribution of the fitness cost. To dissect these effects, one can control for the same growth rate enabling the quantification of changes in PA when WT and mutant strains share an equivalent “physiological state”, i.e., PA_*D*_. To our knowledge, this is the first quantitative description of how RifR mutations modify the global transcriptional program in general, and PA(*μ*) profiles in particular (Fig. 3). That we observe the direct effect of mutations upon promoter activity, PA_*D*_, as an important determinant accentuates the intricate relationship between RNAP activity and fitness. Moreover, in the strain with the most visible fitness costs, there is a significant contribution to changes in PA_*T*_ from the limited availability of global resources.

### General implications

All these results show that decoupling the direct effect is fundamental for a better understanding of the transcriptional reprogramming observed in *rpoB* mutants and its impact on fitness costs. A partially similar approach was used to find a decisive shift in two other *rpoB* mutations whose RNAPs prioritize growth over hedging genes (Utrilla et al. 2016). The authors also compare the genome-wide expression between WT and mutants at a constant growth rate to control for a similar physiological state.

This is a particular example of a more general problem in which the target of a mutation and a phenotype are coupled. Conditions in which a phenotypic change is produced not only by a direct perturbation of a molecular agent, but also by the system-level adaptation to such perturbation are widespread. Some of these systems, but not only, can be found in the context of fitness costs produced by antibiotic resistance mutations when such mutations occur in the molecular target of the antibiotic. Indeed, these perturbations potentially result in complex mutations since antibiotics may impede general cellular functions vital for bacterial growth, for example, DNA replication (quinolones), protein synthesis (macrolides), or transcription (rifamycins) as in our work. But this problem also applies to more specific mutations that also cause genome-scale rewiring. Many open questions remain on whether this rewiring is limited by particular genomic mechanisms, e.g., the possibility of transcriptional compensation (Kafri et al. 2005; Wong and Roth 2005), and thus signifies no fitness costs, or is eventually deleterious, and consequently involves additional costs (Kovács et al. 2020).

Finally, the fact that only a subset of the genes influenced by a complex mutation contributes to fitness appears to subscribe to a model in which extended phenotypic pleiotropy and fitness-relevant modularity coexist (Kinsler et al. 2020). Thus, we notice that many genes –molecular phenotypes– can be affected by these mutations implying extended phenotypic pleiotropy, like that also suggested by genome-wide association studies (Visscher and Yang 2016). However, only a few anticipate fitness hence displaying fitness-relevant modularity like that observed in many laboratory evolution experiments (Tenaillon et al. 2012). We need to continue studying these issues to finally discern how robust function encoded in cells shapes their response to genetic variation.

## MATERIALS AND METHODS

### Computational models of complex mutations

We used the genome-scale metabolic model of *E. coli* iAZ1260 (Feist et al. 2007) together with the Cobrapy toolbox (Ebrahim et al. 2013) to compute the fitness of the WT and mutants in an array of media. We simulated mutations on an enzyme by imposing a limit in the flux of reactions in which it participates. The limit is a fraction of the maximum flux observed across all media in the WT strain and it is fixed for a given mutant during growth in any media. We used minimal media supplemented with one of the 174 carbon sources found in the original study that support growth (Feist et al. 2007). The exchange rate for any carbon source was set equal to that of glucose (8 mmol gDW^−1^ h^−1^). We compute the relative global fitness as the slope of the robust least-squares fit (bisquare method) of the fitness of the mutant relative to the WT. Data points where the mutant is lethal are excluded from the fit. We also used the tool Escher to produce Fig. 1D (King et al. 2015).

### Strains and growth conditions

We used *E. coli* Rel606 as WT, and three mutant derivatives with the following amino acid substitutions in the gene *rpoB*; H526L, S512Y, and Q513P obtained previously through rifampicin resistance. In general, strains were retrieved from −80°C frozen stocks, plated in agar plates with selective media (when necessary), and grown overnight at 37°*C*. Reporter plasmids were extracted from a library (Zaslaver et al. 2006) and purified with the Qiagen Mini-prep kit following the manufacturer’s protocol. Then, each strain was transformed with each reporter plasmid with TSS (Chung et al. 1989). When necessary, selective media for *rpoB* mutants was prepared with rifampicin (100 μg/ml), and for plasmid-bearing strains with kanamycin (50μg/ml). Both antibiotics were used simultaneously when selecting *rpoB* mutants bearing the fluorescent reporter plasmid. All bacterial growth was at 30°C unless otherwise specified. Also, cultures were grown under the shade to prevent rifampicin degradation.

Growth media consisted of M9 minimal media supplemented i) with one of the following carbon sources at 0.5%(w/v): glycerol, sucrose, fructose, and glucose, and ii) either with or without amino acids to a final concentration of 0.2% (w/v), thus making 8 different nutrient conditions in total.

Single colonies were pre-cultured in 1mL of M9 minimal media supplemented with glucose at 0.5%(w/v) for 3h. Then, 96-well flat-bottom plates filled with the corresponding media were inoculated with 20μL of pre-culture to a final volume of 220μL, we then added 30μL of mineral oil to prevent evaporation. Optical density at 600nm, and fluorescence 490/535nm when appropriate, were assayed in a Victor X2 (Perkin Elmer) at 5min intervals with orbital shaking (30s, 1mm) for more than 12h.

### Data processing and promoter activity modeling

First, OD and GFP measurements were corrected for background levels by subtracting the value of blank wells filled with each corresponding growth media. GFP measurements were further corrected by subtracting the autofluorescence produced during the growth of the corresponding strain transformed with the pUA66 promoterless plasmid (Zaslaver et al. 2006). Only then, growth rate time series were computed as the two-point finite differences of log_2_(OD), *μ*(*t*) = ∆ log_2_(OD)/∆*t* (in doublings per hour), and promoter activities were computed as the two-point finite difference in time of fluorescence per OD unit, PA_*pl*_ (*t*) = ∆GFP/∆*t*/OD (in units of GFP/OD/h). Balanced-growth data was computed from the mean time-series measurements of three technical replicates as the average value in a 1h time-window during observable exponential growth.

Promoter activity dependence on growth rate was modelled with a Michaelis-Menten equation as PA(*μ*) = *V*_*m*_ *μ* /(*K*_*m*_ + *μ*) where *V* _*m*_ is the maximum promoter activity and _*m*_ is the growth rate at which PA is half-maximal (Liang et al. 1999). Data from balanced growth was fit to this equation through robust least squares (bisquare) with an upper limit of K_*m*_=3 dbl/h to avoid overfitting linear profiles (Yubero and Poyatos 2020).

## ACKNOWLEDGEMENTS

The authors would like to thank A. Couce and O. Tenaillon for strains. This work was supported by Ph.D. fellowship BES-2016-079127 (P.Y.) and grant PID2019-106116RB-I00 (J.F.P.) from the Spanish Ministerio de Ciencia e Innovación and the European Social Fund.

