## Supplementary Material for "Predicting the fitness costs of complex mutations"

### Supplementary Material
### Predicting the fitness costs of complex mutations

Pablo Yubero and Juan F. Poyatos

March 10, 2021

#### 1 Types of mutations in metabolic models

In this supplementary section, we extend the study of the effect of mutations upon all genes of the metabolic reconstruction iAZ1260 [1]. However, we here restrict mutations to the limiting case of knockouts. Thus, we systematically computed the fitness of all single gene deletions of the metabolic model in minimal media supplemented with one of 174 carbon sources (Methods).

Among all results, we first identify 163 lethal mutations as those with a growth rate $< 0.01$ $h^{-1}$ in all media (Fig. S1A). Second, we find 43 complex mutations with a relative global fitness $|\alpha - 1| > 0.01$ (Fig. S1). Moreover, we repeatedly observe that in some mutations, fitness in a reduced number of media deviates significantly from the general trend. We call these *specific gene-environment interactions* (GxE), and they can be also present in global mutants (Fig. S1AB). For example, we display the environmental fitness cost map of the *atpC* knock-out in Fig. S1B. In fact, *atpC* is part of the ATP biosynthetic process, and apart from displaying a relative global fitness ($\alpha$) it has several specific GxE interactions.

Figure S1C shows the genes that we have identified as complex mutations and the magnitude of their associated relative global fitness (Methods), which ranged from 20% to a mild 105%. A global fitness $> 1$ should be considered with caution as they are associated to a larger increase in growth rate between two carbon sources compared to the wild type.

Finally, we tested the idea that genes whose knock out produces smaller relative global fitness (larger costs) could be associated to larger flux changes due to the mutation. We measured flux changes as the euclidean distance between the flux vectors of the WT and the mutant strain in all media that permitted growth, and found a clear and strong negative linear correlation ($\rho = -0.61$ p<0.001, Fig. S1D). In addition, we tested a similar hypothesis against the number of reactions in which each enzyme participates. We find, that the more reactions are impeded by the knock out of an enzyme, the smaller the relative global fitness, although with a small correlation coefficient ($\rho = -0.24$ p<0.001, Fig. S1E). Overall, this demonstrates that in this computational model, complex mutations reconfigure to a larger extent the metabolism than "simple" mutations.

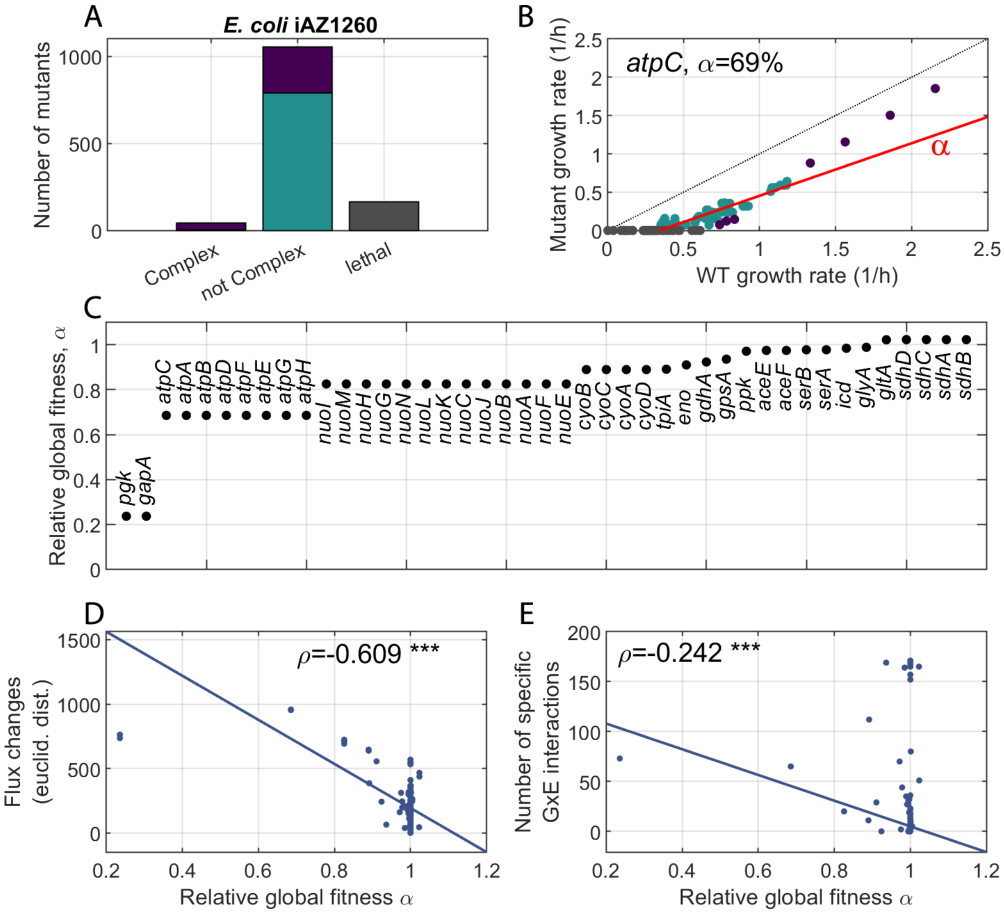


**Figure S1. Complex mutations in the whole-genome metabolic model iAZ1260 of *E. coli*.** (**A**) Distribution of the type of mutations found across all single-gene deletions. We find 43 complex mutations, 1054 "simple" mutations and 163 lethal mutations across all growth media (gray). We also display the number of mutants that display none (turquoise), and at least a specific GxE interaction (purple). (**B**) Environmental fitness cost map of *atpC* across all 174 growth media. We find some lethal GxE interactions (gray), and a group of points (turquoise) that follow a general trend (red solid line). We quantify this trend by its slope $\alpha$ which we call the *global relative fitness*. There are also some points that deviate significantly from this trend and that are automatically considered specific GxE interactions (purple, Methods). Finally, we also show the wild type reference (diagonal black dotted line). (**C**) Global relative fitness of all complex mutations identified in the single-gene deletion collection. (**D**) The relative global fitness of a mutant correlates negatively with the flux changes produced by the mutation ($\rho$ linear correlation coefficient, *** $p<0.001$). (**E**) Similarly but to a lesser extent, the relative global fitness of a mutant correlates negatively with the number of reactions in which the mutated gene participates ($\rho$ linear correlation coefficient, *** $p<0.001$).

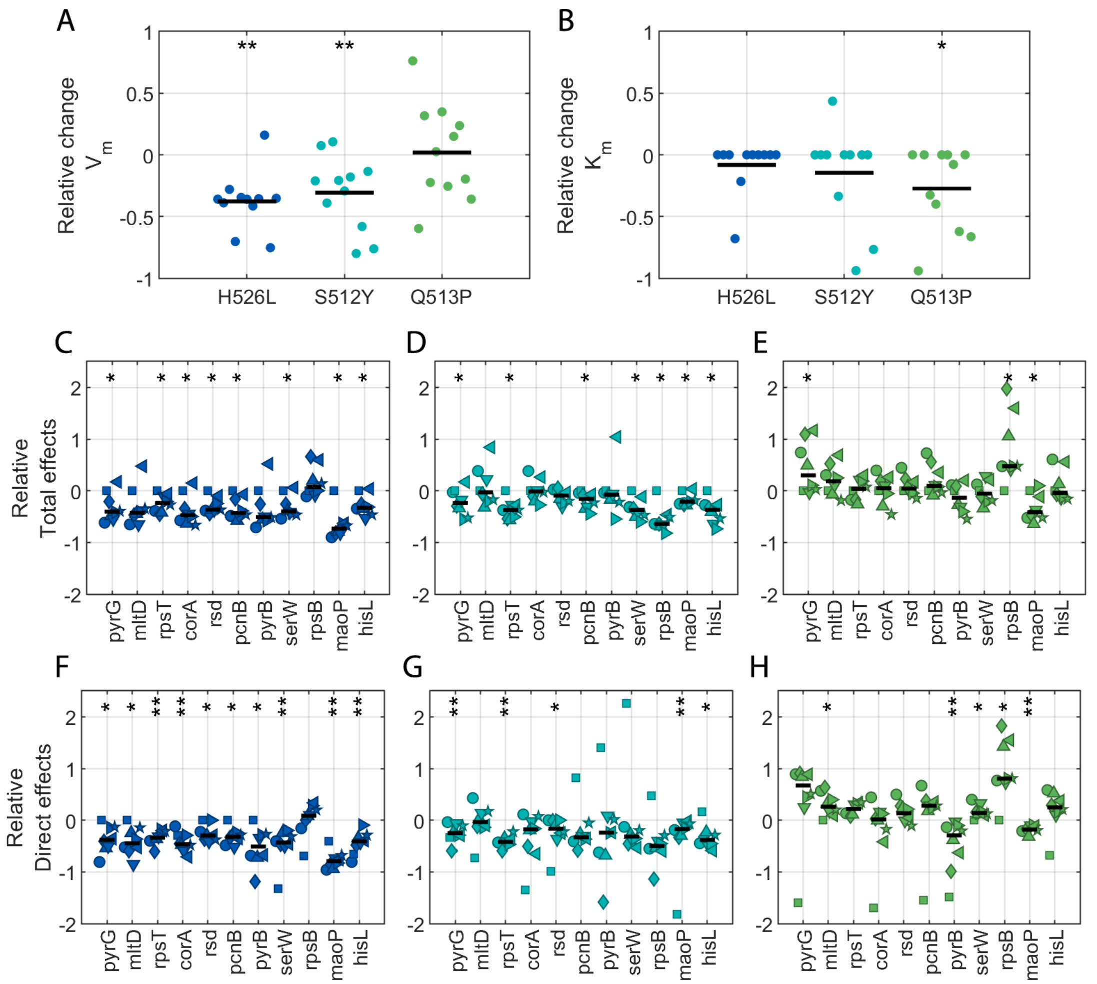


**Figure S2. Mutations in *rpoB* modify the global transcriptional program.** Relative changes of the fitting parameters (**A**) $V_m$ and (**B**) $K_m$ to the PA($\mu$) profiles of each promoter (dots) and each strain (colors, see Methods and Fig.3). (**C-E**) Total, and (**F-H**) direct effects, relative to the wild type of all promoters. We show data for all growth media (symbols, as in Fig.2) and its median (black horizontal bars). In all panels, we tested for a general tendency of increased or decreased value across growth media with a two-side Wilcoxon sign test for medians (* $p<0.05$, ** $p<0.01$).

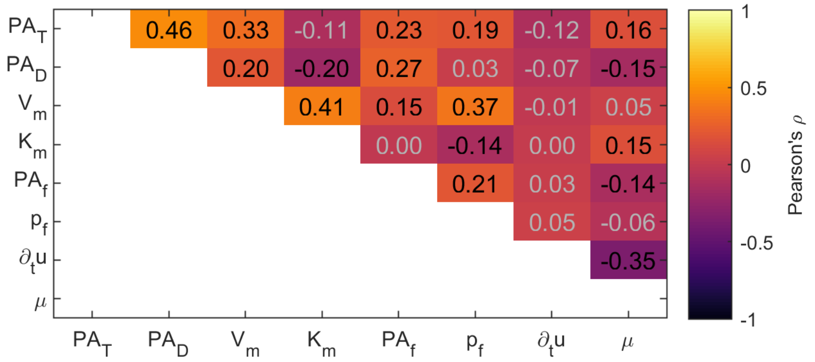


**Figure S3. Cross-correlations of all variables of the statistical model , including fitness.** We show the linear cross correlations among all fitness predictors: total effects on PA ($PA_T$), direct effects on PA ($PA_D$), maximum PA ($V_m$), growth-rate at which PA is half-maximal ($K_m$), PA in stationary phase ($PA_f$), GFP concentration in stationary phase ($p_f$), deceleration rate at the exit of balanced growth ($\partial_t \mu$) and fitness ($\mu$). Data was aggregated across promoters and growth media, and all values are relative to the wild type, as in the statistical model. Values that are significant, with p<0.05, are shown in black.

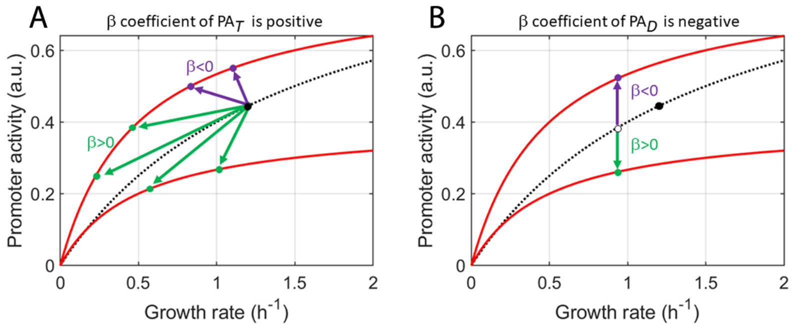


**Figure S4. Impact on PA($\mu$) of the coefficient sign of $PA_T$ and $PA_D$** In both panels we show the PA($\mu$) profiles associated to a hypothetical promoter in the wild type (black dotted line) and two mutants (red solid lines) to assess the impact of the sign of the coefficient, $\beta$, obtained in the statistical model. (**A**) In the case of $PA_T$, the sign of $\beta$ is associated to the mapping between PA and growth rate $\mu$ in a fixed growth media between the wild type (black circle) and mutant (colored circles). That the sign is positive, delimits such possible mapping to those similar to the green arrows, where a decrease in growth rate (fitness costs) produce a decrease in $PA_T$. (**B**) In the case of $PA_D$, the sign of the coefficient is negative. Thus, fitness costs are associated to the over expression of the promoter (purple arrow) rather than under expression (green arrow) from the expected PA of the wild type at the growth rate of the mutant (black empty circle). Both results, when combined give a general view of how PA($\mu$) profiles change due to mutations in *rpoB*.

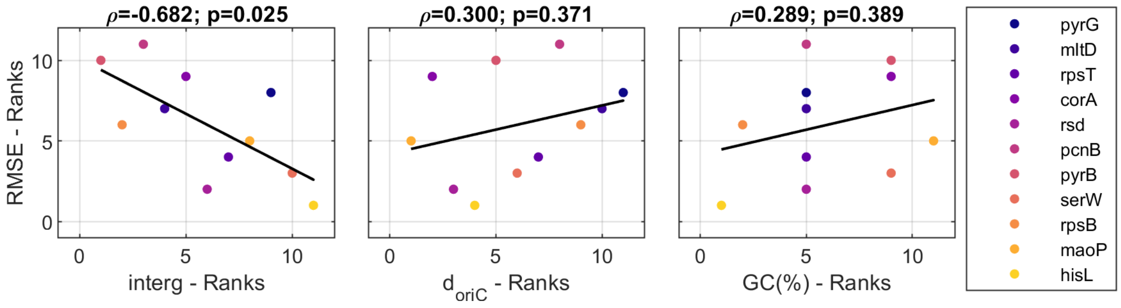


**Figure S5. Genes that perform better at fitness prediction have a larger intergenic distance.** To test whether the good performance in anticipating fitness costs of some genes can be derived from molecular attributes we display the rank correlations between the goodness of fitness predictions (RMSE, see Fig.5) and different gene-associated parameters. We find that the goodness of fit (RMSE) of the selected genes correlates significantly with the intergenic distance (Spearman's $\rho = -0.68$), as opposed to the distance to *oriC* ($d_{oriC}$) or the GC content of the promoter region. We also show rank correlation $\rho$ and their p value. Finally, neither gene essentiality nor regulation from different sigma factors seem determinants of good predictors (not shown).

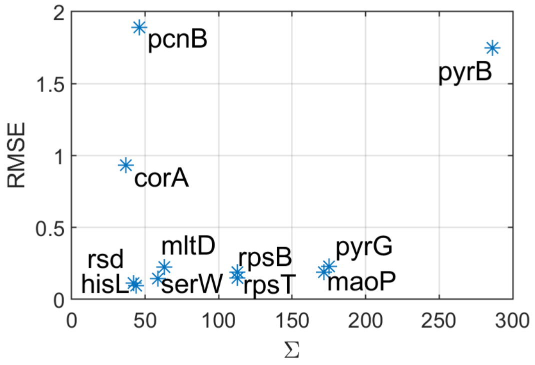


**Figure S6. Genes that are better predictors lie in an optimal sensitivity to the mutation.** We quantified the sensitivity of a gene to a mutation as the sum of the squared relative change wrt the wild type of all predictors ($\Sigma$). That we observe a U-shaped tendency suggests that genes that have either very large or low sensitivity to a mutation are worse predictors (large RMSE) of fitness costs that those having intermediate values.

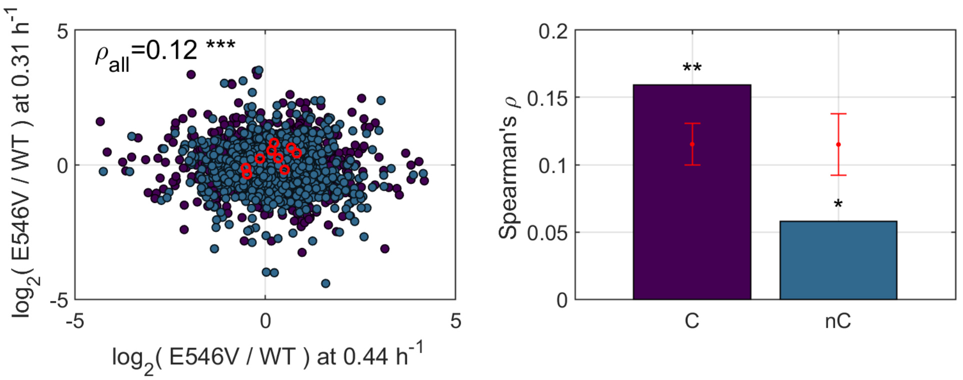


**Figure S7. Transcriptomic changes in *rpoB* mutant E546V when controlling for growth rate.** (**A**) Fold change of the number of transcripts of the E546V mutant growing at two different rates correlate only slightly (Spearman's $\rho_{all} = 0.12$). Each point corresponds to a gene, and red circles indicate the genes used in this study, data for *hisL* and *serW* are not available. (**B**) Rank correlations of the subset of constitutive (C, purple) and regulated (nC, blue) genes. Red dots and error bars represent the mean and one standard deviation of a $10^4$ permutation test. Constitutive genes correlate significantly more than regulated genes (* p<0.05; **p<0.01, two-tailed).
